# WELPCR: Low-cost polymerase chain reaction (PCR) thermal cycler for nucleic acid amplification and sensing

**DOI:** 10.1101/2023.09.11.557292

**Authors:** Poulami Mandal, Vivekanand Dhakane, Avani Kulkarni, Siddharth Tallur

**Author notes:** equal contribution (Poulami Mandal, Vivekanand Dhakane).

## Abstract

The COVID-19 pandemic highlighted the need for sensitive and cost-effective diagnostics. The gold standard test to diagnose infectious diseases such as COVID-19 is quantitative polymerase chain reaction (qPCR). Although there have been multiple advances in nucleic acid testing, PCR thermocyclers are expensive and typically not affordable for offering hands-on training to PCR in instructional laboratory courses in undergraduate colleges. We introduce a low-cost, portable, and automated real-time PCR device designed at the Wadhwani Electronics Lab (WEL) at IIT Bombay, christened WELPCR. WELPCR is an open-source design, and capable of reverse transcription PCR (RT-PCR) for complementary DNA synthesis, as well as end-point PCR of small amount of DNA in a test sample. The WELPCR design enables ramp rate up to 2 °C*/*s, weighs less than 2 kg, and is easy to assemble. The built-in LED display and keypad provides an intuitive and seamless user interface to configure and set up WELPCR. With bill of materials (BOM) cost less than INR Rs. 10 000, WELPCR is an attractive solution for bringing PCR technology to instructional and research laboratories in a cost-effective manner.

## I. Introduction

Polymerase chain reaction (PCR) amplifies DNA in a rapid and highly specific manner, allowing generation of millions to billions of copies of target DNA. At the heart of a PCR machine is a thermal cycler, that rapidly switches across three different temperatures for denaturation, annealing and extension of the amplicons. It is essential to maintain these temperatures accurately with fast transition between these temperatures to have a shorter overall turnaround time of the reaction. To develop a PCR device, it is essential to consider the need for accurate heating and temperature sensing requirements in such a way that the device is low-cost and not bulky. While there are several highly sophisticated and large PCR machines for high throughput sample analysis available on the market, these instruments are typically very expensive and cost well in excess of USD $2000. There are also smaller, less expensive designs such as miniPCR® (approximately USD $500) and similarly-priced open-source designs such as OpenPCR [1], that make this technology more accessible. However, the price point is relatively high for a large number of instructional and research laboratories at educational institutes in low and middle income countries (LMICs). We present WELPCR, a low-cost and portable PCR thermal cycler developed at the Wadhwani Electronics Lab (WEL), IIT Bombay. The bill of materials (BOM) cost for assembling one unit of WELPCR is less than INR Rs. 10 000 (approximately USD $120), which makes it a great resource for educational and research use in laboratories with limited resources. The WELPCR design is open-source and made available for developers interested in assembling their own units [2].

## II. Design

The WELPCR design is inspired by the OpenPCR design [1]. WELPCR is a standalone device and does not require connection to a computer or a mobile device for configuration and operation. Figure 1 shows the block diagram of the design. Detailed description of the various elements, along with detailed assembly and operation instructions are provided in the project documentation on GitHub [2]. A brief description of various blocks in the system is presented in this section below. The mechanical assembly includes manufacturing the tube holder, lid bracket and sheet metal parts. 3D parts were machined using CNC milling at the Machines and Tools Lab, Department of Mechanical Engineering, IIT Bombay. The back panel is manufactured using laser-cut aluminum sheet metal and the rest of the mechanical body is assembled using laser-cut acrylic sheets. Acrylic casing faces are usually fixed with an adhesive. However, keeping in mind the need to disassemble the casing for servicing and repairs, we have used L-bracket with nut-bolt to connect two perpendicular panels. For maintenance, we need to open only one face of the casing, and hence the front face is fixed using a hinge to allow it to swivel like a door. Figure 2 shows the 3D CAD drawing photograph of a fully assembled WELPCR unit.

**Fig. 1.**
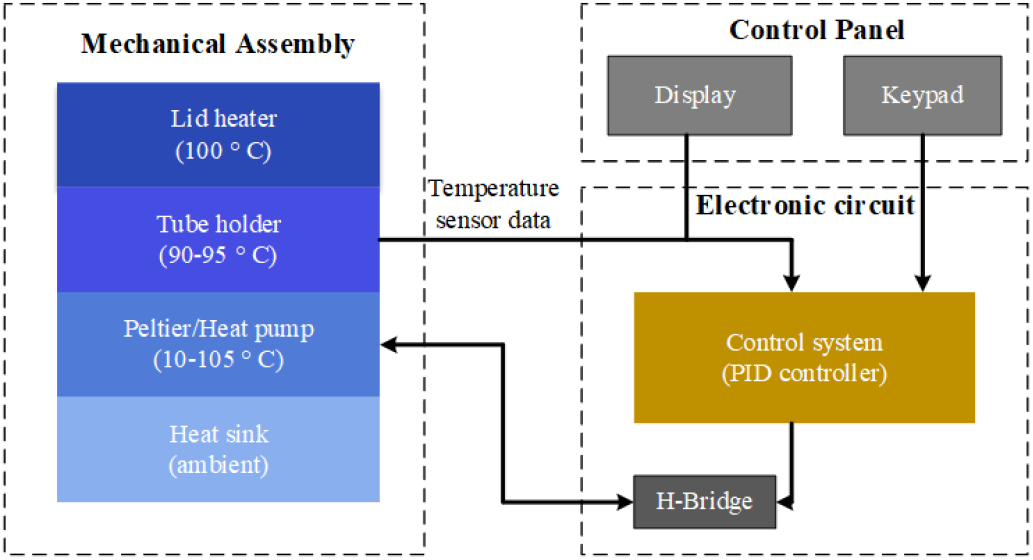
Block diagram of WELPCR

**Fig. 2.**
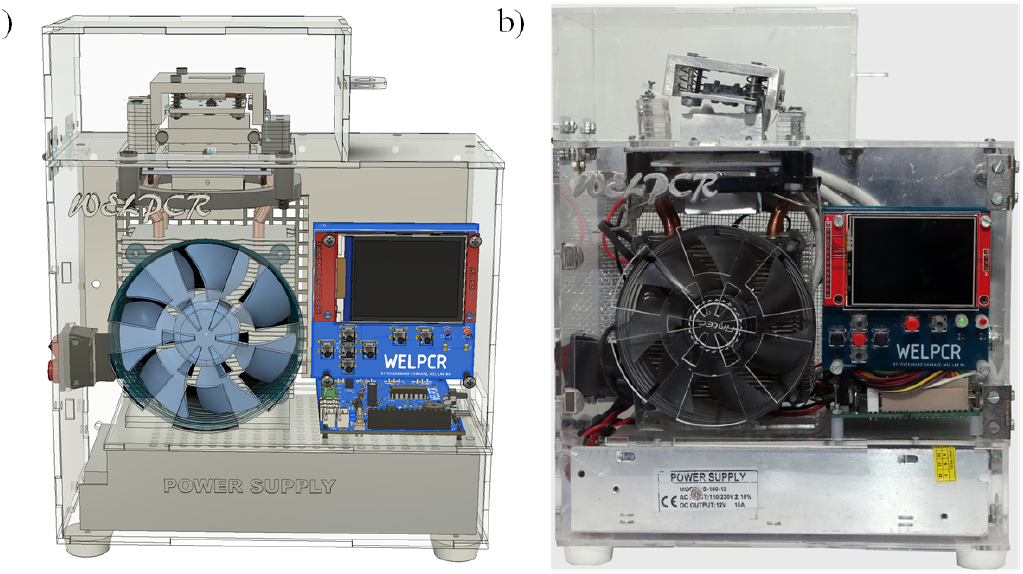
(a) WELPCR designed in CAD tool (Autodesk® Fusion 360) (b) Photograph of fully-assembled WELPCR. after manufacturing

### A. Mechanical design

Various components of the mechanical design shown in Figure 3 are described in brief below:

**Fig. 3.**
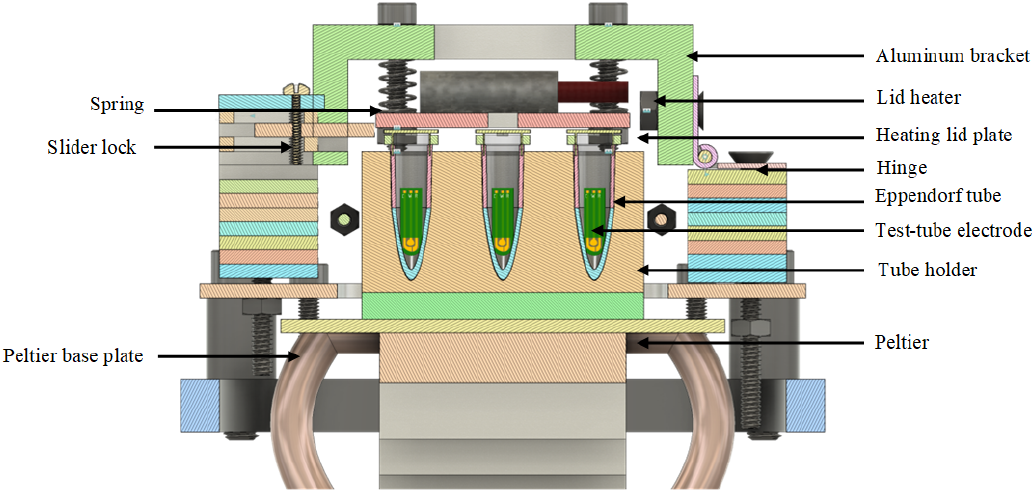
Cross-sectional view of sample holder and thermal cycler mechanical assembly.

**Fig. 4.**
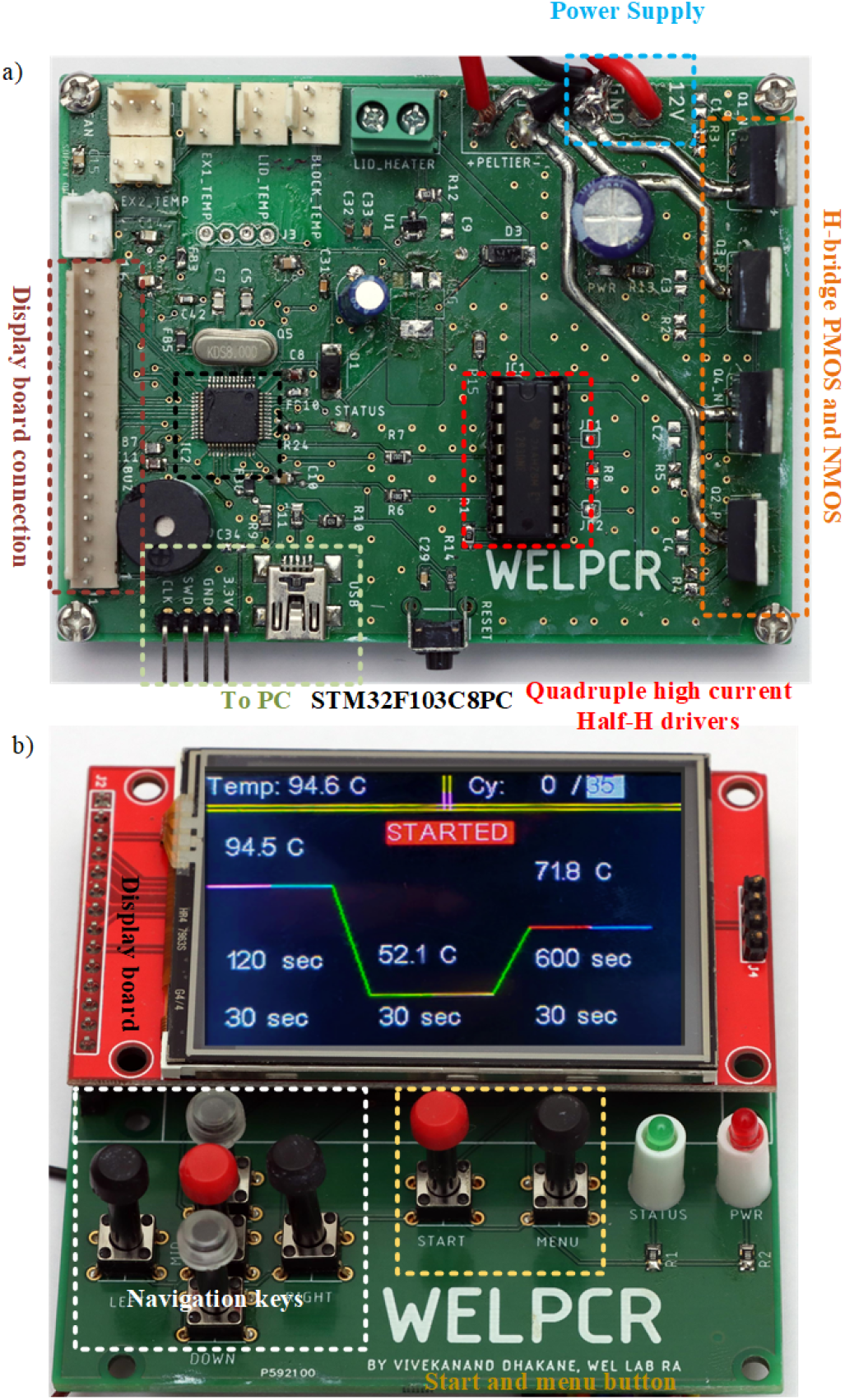
Top-view picture of the PCR circuit board with labels corresponding to schematic components a) PCR motherboard b) PCR display board

#### a) Tube holder

The tube holder contains 3 wells for inserting 0.2 ml Eppendorf tubes that will contain the sample. This block is heated using a Peltier cell.

#### b) Peltier cell

A Peltier cell is a heat pump that can pump heat from one side to the other side depending on the direction of current flow. We can heat and cool the tube holder by simply inverting the current flow direction using an Hbridge.

#### c) Lid heater and heating lid plate

Lid heater is a ceramic 12 V 40 Wceramic cartridge heater, commonly used in 3D printers. This heating element is used to heat the lid of the tube holder, to avoid condensation of sample and PCR reagents at the top of the tube while the PCR reaction is running. The solid square bracket ‘[’ shaped lid is pressed into contact with the Eppendorf tube cap using a spring-loaded assembly. heating plate.

#### d) Fan

A CPU cooling fan is used for controlling air flow to sink (while decreasing the temperature) and source (while increasing the temperature) heat to aid the temperature cycling.

#### e) Peltier base plate

Peltier base plate is used as an interface between the CPU cooling fan and the Peltier cell, for efficient heat transfer.

### B. Electronics design

The main sub-systems in the electronics module include the heating element driver, temperature controller and user interface. The core temperature cycling functionality is realized using a Peltier cell, driven with a custom-developed 15 A Hbridge driver for bi-directional current (heating and cooling). The H-bridge circuit has two PMOS (IRF9540) and two NMOS (IRF540) transistors. A proportional-integral-derivative (PID) controller is designed to control and quickly stabilize the temperature of the tube holder for each amplification stage (denaturation, annealing and extension). As PCR is extremely temperature-sensitive and primers may be damaged because of overshoot in the temperature, we need a critically damped PID controller. Temperature of the tube holder blow was measured with a thermistor and used to set the duration of the pulse width modulation (PWM) signal for the Peltier cell driver:

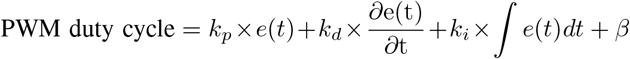

where

*k*_*p*_ = proportional gain

*k*_*d*_ = derivative gain

*k*_*i*_ = integral gain

e(t) = error signal (difference between set point and measured temperature)

*β* = bias

The gains are empirically determined.

## III. Experimental results

### A. PCR cycle

PCR cycles through different temperatures with different time duration. The first step is denaturation which requires (90–95) ^*°*^C [3]. It causes denaturation of the DNA i.e., DNA is broken into single-stranded DNA template. The second step is annealing, which requires (50–60) ^*°*^C. In this step, primers are annealed to each of the single-stranded DNA templates. The third step is extension, which requires 72 ^*°*^C. In this step, new DNA strand complementary to the DNA template strand is synthesized. Each cycle doubles the number of DNA copies in the sample. This cycle is repeated *N* number of times, resulting in increase in DNA concentration by a factor of 2^*N*^ . Figure 5 shows the temperature profile for two cycles of thermal cycling measured on WELPCR. The blue curve shows the temperature profile of the tube holder, which is directly attached to the Peltier cell. The orange curve shows the temperature profile of the sample inside the Eppendorf tube, measured with a thermistor. The sample temperature is observed to be usually lesser than the tube holder temperature by (1–2) ^*°*^C, because the sensor measuring the temperature of the holder is closer to the heating element, and due to heat loss at the Eppendorf tube-tube holder interface. Heat loss at this interface is minimized using thermal conducting paste. To measure the temperature in tube holder and sample tube, we used a 1 mm, 4.7 k? thermistor. The temperature sensor needs to be as small as possible because the sample volume is 20 *µ*l, which occupies only (2–3) mm depth in the test tube. Further, a big thermistor also adds a thermal load, and results in heat loss as well as erroneous temperature readings because of inadequate contact with the small amount of sample.

**Fig. 5.**
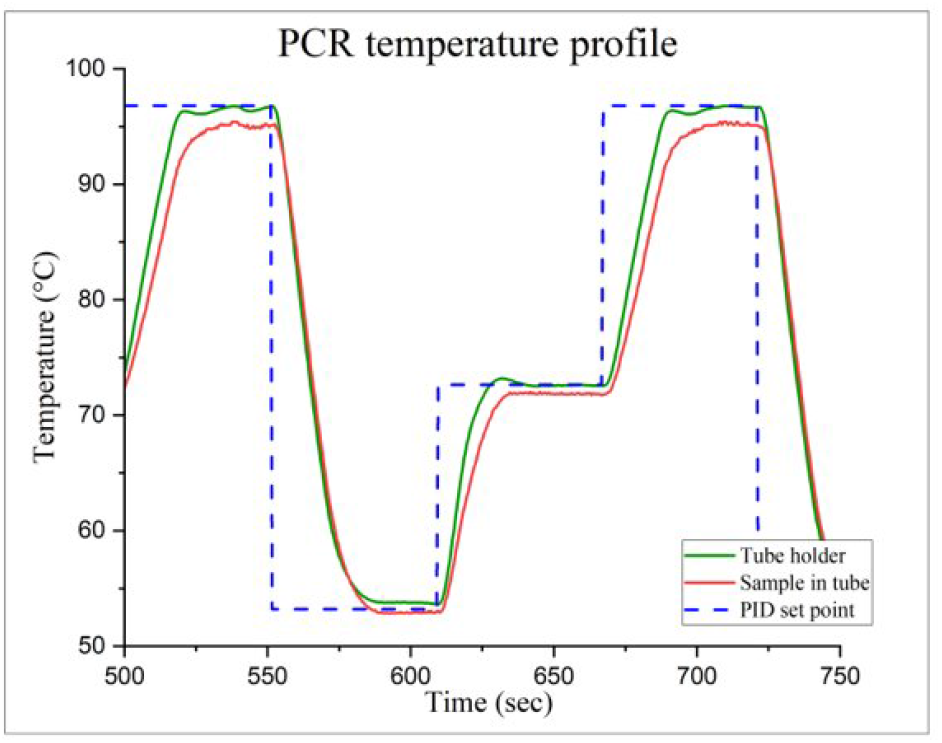
WELPCR temperature profile

### B. Sample preparation

For demonstrating the utility of WELPCR, we amplified nucleic acid from bacteriophage phi6, an enveloped doublestranded RNA (dsRNA) virus that infects *Pseudomonas syringae* (non-pathogenic BSL-1 strain). This virus serves as a surrogate for highly pathogenic enveloped viruses such as SARS-CoV-2, ebola and influenza [4], [5]. Using GenElute™ Universal Total RNA purification kit (Sigma-Aldrich), RNA extraction was carried out as per the manufacturer’s protocol. Extracted RNA was quantified by determining the absorbance at 260 nm using a UV/Vis spectrophotometer (Thermo Scientific Multiskan GO) [5]. The RNA extracted was used as template for further PCR reactions. Detailed sample preparation procedure is reported in our earlier work [5]. Amplification was carried out parallelly in WELPCR and a commercially available thermocycler (Bio-Rad T100™ Thermal Cycler) to compare the performance.

### C. cDNA synthesis and amplification of target

cDNA was synthesized on the both instruments using the iScript cDNA synthesis kit (Bio-Rad Laboratories) [6] as per the manufacturer’s protocol. Briefly, it consists of three steps: priming at 25 ^*°*^C for 5 min, reverse transcription at 46 ^*°*^C for 20 min and inactivation of reverse trancriptase enzyme at 95 ^*°*^C for 1 min [5]. Synthesised cDNA was used as a template for amplifying a 503 bp long fragment. A 20 *µ*l reaction was set up using 500 nM forward and reverse primers and 2*X* PCR Master Mix (Thermo Fisher Scientific). Following are the primer sequences

Forward: 5’-GAACCATATGACTTTGTACCTGGTCC-3’

Reverse: 5’-CAACGAATTCTCAGGCGCTTACCTCATC-3’

Amplification was performed with initial denaturation stage at 95 ^*°*^C for 2 min followed by 35 cycles of thermal cycling (95 ^*°*^C for 30 s, 52.8 ^*°*^C for 30 s and 72 ^*°*^C for 30 s). Final stage extension was carried out at 72 ^*°*^C for 10 min. Time required for completing the amplication was 106 min on BioRad T100™ Thermal Cycler and 114 min on WELPCR.

### D. Quantification of PCR product

The amplified PCR product were loaded on to 1 % agarose gel and visualized using GelDoc Go Gel Imaging System (BioRad Laboratories). To compare the performance of WELPCR with Bio-Rad T100™ Thermal Cycler in terms of amplification, the obtained PCR product was purified using GeneJET Gel Extraction Kit (Thermo Fisher Scientific) and purified DNA was eluted in 30 µl of nuclease free water. This purified DNA was quantified using NanoDrop™ Lite Spectrophotometer.

### E. Results and Discussion

#### a) Analysis of amplified product

In order to compare the efficiency of the presented thermal cycler with commercially available, cDNA synthesis was setup on WELPCR and BioRad T100™ Thermal Cycler. Using the synthesized cDNA from respective thermal cyclers, a PCR reaction was set up, and the gel image obtained after electrophoresis is shown in Figure 6. Visual inspection of the gel shows that band intensity for WELPCR is comparable to that of Bio-Rad T100™ Thermal Cycler. This suggests that the amplification performance of WELPCR is similar to that of Bio-Rad T100™ Thermal Cycler.

**Fig. 6.**
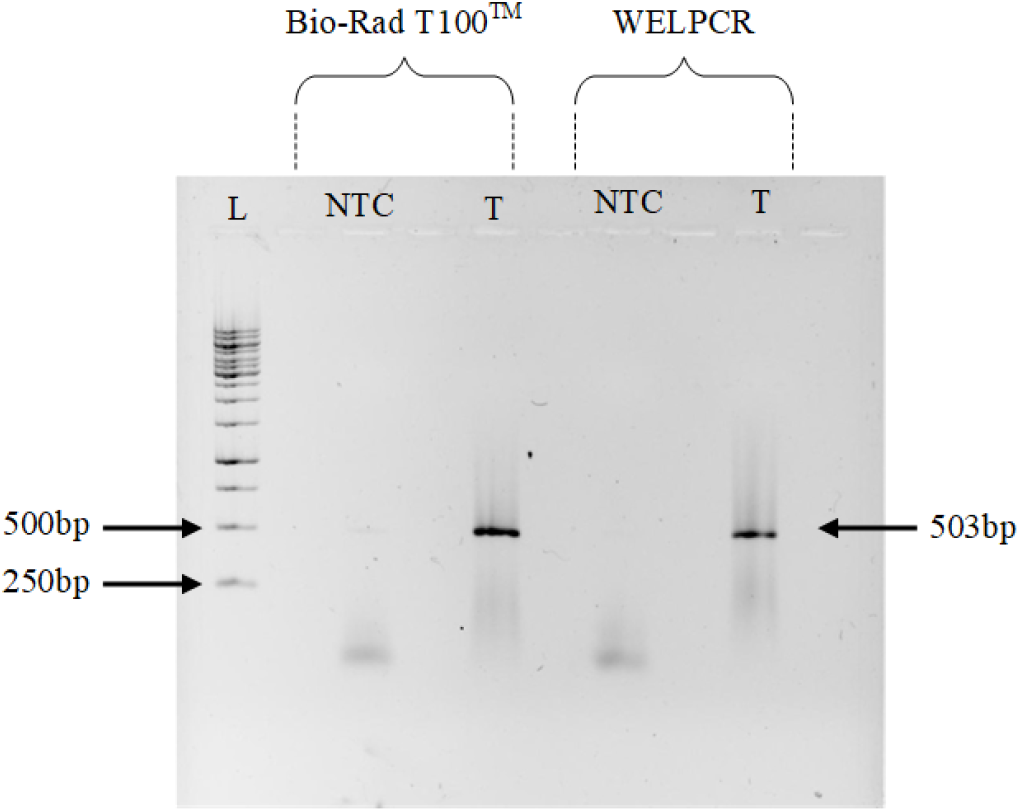
Agarose gel electrophoresis image showing amplified phi6 cDNA using both Bio-Rad T100™ Thermal Cycler and WELPCR, L: Ladder, NTC: No template control, T: Test sample (phi6)

#### b) Quantitative analysis

To determine the amount of nucleic acid after amplification and compare it with BioRad MJ MiniTM Personal thermal Cycler, the PCR product was purified and 2 µl of the product was used to determine the concentration. The concentration as determined using NanoDrop™ Lite Spectrophotometer was determined to be 20.1 ng*/*µl for Bio-Rad T100™ and 16.5 ng*/*µl for WELPCR.

## IV. Conclusion

In summary, we present a low-cost, portable and easy-to-assemble and operate PCR thermal cycler, WELPCR. The performance specifications of WELPCR and comparison with other low-cost options is shown in Table I. The utility of the WELPCR was shown through reverse transcription and amplification of nucleic acid from bacteriophage phi6, a popular surrogate virus for SARS-CoV-2. All these attributes make WELPCR a suitable thermal cycler for resource-constrained laboratories for teaching and research use. To enable easy adoption of the technology, WELPCR design is open-sourced and all design resources and assembly and testing instructions are provided on GitHub [2].

**TABLE I.** WELPCR comparison with other low-cost PCR machines.

| Specification | WELPCR | OpenPCR | miniPCR® mini8 |
| --- | --- | --- | --- |
| Maximum ramp rate | 2 °C/s | 1 °C/s | 2.4 °C/s |
| Reverse transcription | Yes | No | No |
| No. of wells | 3 (extendable to 9) | 16 | 8 |
| Temperature range | (10–105) °C | (20–105) °C | Ambient–99 °C |
| User Interface | Keys, touchscreen Display | Computer software | Mobile app |
| Weight | 1.9 kg | NA | 0.45 kg |
| Cost (INR) | Rs. 10,000 (BOM) | Rs. 38,500 (BOM) | Rs. 75,000 + shipping and taxes |

## Acknowledgments

The authors thank all staff at the Wadhwani Electronics Lab (WEL), IIT Bombay, for their support.

